# VECTRON™ T500, a new broflanilide insecticide for indoor residual spraying, provides prolonged control of pyrethroid-resistant malaria vectors

**DOI:** 10.1101/2022.10.12.511866

**Authors:** Renaud Govoetchan, Augustin Fongnikin, Thomas Syme, Graham Small, Martial Gbegbo, Damien Todjinou, Mark Rowland, Derric Nimmo, Germain Gil Padonou, Corine Ngufor

## Abstract

**Background:** Broflanilide is a newly discovered insecticide with a novel mode of action targeting insect γ-aminobutyric acid receptors. The efficacy of VECTRON™ T500, a wettable powder formulation of broflanilide, was assessed for IRS against wild pyrethroid-resistant malaria vectors in experimental huts in Benin.

**Methods:** VECTRON™ T500 was evaluated at 100 mg/m² in mud and cement-walled experimental huts against wild pyrethroid-resistant *Anopheles gambiae sensu lato* (*s*.*l*.) in Covè, southern Benin, over 18 months. A direct comparison was made with Actellic® 300CS, a WHO-recommended micro-encapsulated formulation of pirimiphos-methyl, applied at 1000 mg/m². The vector population at Covè was investigated for susceptibility to broflanilide and other classes of insecticides used for vector control. Monthly wall cone bioassays were performed to assess the residual efficacy of VECTRON™ T500 using insecticide susceptible *An. gambiae* Kisumu and pyrethroid-resistant *An. gambiae s*.*l*. Covè strains. The study complied with OECD principles of good laboratory practice.

**Results:** The vector population at Covè was resistant to pyrethroids and organochlorines but susceptible to broflanilide and pirimiphos-methyl. A total of 23,171 free-flying wild pyrethroid-resistant female *An. gambiae s*.*l*. were collected in the experimental huts over 12 months. VECTRON™ T500 induced 56%-60% mortality in wild vector mosquitoes in both cement and mud-walled huts. Mortality with VECTRON™ T500 was 62%-73% in the first three months and remained >50% for 9 months on both substrate-types. By comparison, mortality with Actellic® 300CS was very high in the first three months (72%-95%) but declined sharply to <40% after 4 months. Using a non-inferiority margin defined by the World Health Organization, overall mortality achieved with VECTRON™ T500 was non-inferior to that observed in huts treated with Actellic® 300CS with both cement and mud wall substrates. Monthly *in situ* wall cone bioassay mortality with VECTRON™ T500 also remained over 80 % for 18 months but dropped below 80% with Actellic® 300CS at 6-7 months post spraying.

**Conclusion:** VECTRON™ T500 shows potential to provide substantial and prolonged control of malaria transmitted by pyrethroid-resistant mosquito vectors when applied for IRS. Its addition to the current list of WHO-approved IRS insecticides will provide a suitable option to facilitate rotation of IRS products with different modes of action.

## Background

Indoor residual spraying (IRS) has contributed substantially to the overall reduction in malaria burden seen across Africa over the last two decades [1]. It is a core vector control intervention that greatly reduces indoor adult mosquito vector density and longevity, for several months following its application [2, 3]. This reduction in both biting rate and survival of vectors, considerably lowers indoor malaria transmission risk. Despite the undoubted effectiveness of IRS demonstrated in many transmission settings [4-7], the high prevalence of resistance in *Anopheles* vectors to the limited number of classes of insecticides available for this intervention [8], is of serious concern. Pyrethroid resistance is now widespread and increasing in intensity in major malaria vectors across Africa while resistance to the three other insecticide classes commonly used in IRS up to 2016 (organophosphates, carbamates and DDT), has been confirmed across malaria endemic countries in the five World Health Organization (WHO) regions [8]. The threats of insecticide resistance in malaria vectors and the lack of suitable cost-effective alternative IRS insecticides have contributed to an overall decrease in the number of structures sprayed [9] and a reduction in the percentage of the population at risk protected by IRS, from 5.8% in 2010 to 2.6% in 2020 [10]. This has been followed by disturbing reports of malaria resurgence following IRS withdrawal in many countries [11-13].

There is an urgent need for new effective and long-lasting alternative chemistries for indoor residual spraying [14]. New IRS insecticide formulations containing clothianidin, a neonicotinoid insecticide, were recently added to the WHO list of vector control products [15]. However, to help increase capacity to manage insecticide resistance in malaria vectors through the rotation of insecticides for IRS as recommended by the WHO [14], multiple insecticides with new modes of action need to be developed. Mitsui Chemicals Agro, Inc. (MCAG) has developed a new active ingredient, broflanilide, a meta-diamide insecticide with a novel mode of action, classified as a GABA-gated chloride channel allosteric modulator [16-18]. Earlier studies demonstrated its effectiveness against a wide range of insect pests [19-21] and since 2020, it has been marketed worldwide for control of agricultural pests [22]. Based on its novel mode of action, broflanilide is also being developed for public health use [17, 23]. Preliminary laboratory bioassays demonstrated the efficacy and residual activity of early formulations of the insecticide on various household substrates against malaria vector mosquitoes and the absence of cross-resistance to most insecticides currently used for vector control [24]. More recently, a wettable powder formulation of this insecticide developed for indoor residual spraying (VECTRON™ T500) demonstrated prolonged efficacy on mud and cement wall substrates in laboratory bioassays and in preliminary animal baited experimental hut evaluations against wild-free flying malaria vector mosquitoes in Benin [23] and Tanzania [25]. Based on the findings from these trials and on toxicity studies performed with the insecticide, an application rate of 100mg/m^2^ was selected as a suitable dose for indoor residual spraying with VECTRON™ T500.

To be added to the WHO list of vector control products, a new IRS insecticide should ideally demonstrate non-inferiority to a WHO-recommended IRS insecticide in semi-field experimental hut studies [26]. Such studies should preferably be performed following the OECD principles of good laboratory practice (GLP) [27]. To generate efficacy data as part of a WHO PQT/VCP dossier submission for the assessment of VECTRON™ T500 for indoor residual spraying against malaria vectors, an experimental hut trial in human occupied experimental huts was conducted to assess the efficacy and residual activity of VECTRON™ T500 applied at 100 mg AI/m^2^ against pyrethroid-resistant *Anopheles gambiae sensu lato* (*s*.*l*.) in Covè, southern Benin. VECTRON™ T500 was also investigated for its non-inferiority to Actellic® 300CS, a WHO PQT/VCP listed IRS insecticide formulation of the organophosphate insecticide pirimiphos-methyl. The study was conducted following OECD principles of GLP at the GLP certified CREC/LSHTM Collaborative Research Programme in Benin.

## Methods

### Study site and experimental huts

The experimental hut trial was conducted at the CREC/LSHTM experimental hut station in Covè, Southern Benin (7.21’N-2.34’E). The hut site is located in an irrigated valley producing rice throughout most of the year and providing suitable breeding habitats for mosquitoes. The rainy season extends from March to October and the dry season from November to February. The vector population consists of both *Anopheles coluzzii* and *An. gambiae sensu stricto* (*s*.*s*.) with the latter occurring at lower frequencies (∼23%) and mostly in the dry season [28]. The vector population is highly resistant to pyrethroids. Molecular genotyping and microarray studies have demonstrated a high frequency of the L1014F allele (> 90%) and overexpression of the cytochrome P450s CYP6P3, associated with pyrethroid detoxification [28].

The hut trial ran for 18 months between November 2019 and May 2021 in 8 experimental huts of West African design [29]. Wild mosquitoes were allowed to enter in the experimental huts for first 12 months while in situ wall cone bioassays were performed for 18 months. The experimental huts are made from concrete bricks with a corrugated iron roof. Inner walls and ceiling were plastered with either concrete or mud and the ceilings were made of the same materials as the walls (mud or cement). Both wall substrates were prepared in line with local practices; 1:3 ratio of cement and sand for concrete walls and 2:3 ratio of mud and sand for mud walls. To improve the durability of the mud wall plaster in line with local practice, a small amount of cement (10%) was added to the mud sand mix. To prevent any contamination from previous trials, hut walls were refurbished by replastering and the substrates were allowed to cure for 1 month before the evaluation. Each hut was built on a concrete plinth surrounded by a water-filled moat to prevent the entry of scavenging ants and had a wooden framed veranda trap to capture exiting mosquitoes. Mosquito entry occurred via four window slits each measuring 1 cm in width and situated on three sides of the hut.

### Susceptibility of wild vector population to broflanilide

Bioassays were conducted to assess the susceptibility of the Covè vector population to broflanilide, organochlorines, pyrethroids, organophosphates and carbamates during the hut trial. Bottle bioassay were used to investigate to susceptibility to broflanilide at a pre-determined diagnostic dose of 6 µg/bottle using 800 ppm Mero® (Bayer CropScience, Germany) as adjuvant while WHO cylinder bioassays were used to investigate resistance to organochlorines (DDT 4%, dieldrin 4%) pyrethroids (permethrin 0.75%, alpha-cypermethrin 0.05%, and deltamethrin 0.05%), carbamates (bendiocarb 0.1%) and organophosphates (malathion 5% and pirimiphos-methyl 0.25%) with filter papers obtained from Universiti Sains Malaysia [30]. For both types of bioassays, 100 unfed wild *An. gambiae s*.*l*. Covè adult mosquitoes that emerged from larvae collected from breeding sites within the experimental hut site were exposed in cohorts of 25 mosquitoes per bottle or tube for 1 hour and delayed mortality was recorded 72h later for broflanilide (due to the slower acting mode of action of this insecticide) and 24 h later for the other classes of insecticides. Control mosquitoes were exposed to untreated papers or to bottles coated with acetone and 800 ppm Mero®. Tests were also performed with the insecticide-susceptible *An. gambiae s*.*s*. Kisumu strain for comparison.

### Experimental hut treatments

A total of six (6) treatments were evaluated in eight (8) experimental huts (Table 1). The performance of VECTRON™ T500 was assessed on concrete and mud walled huts at the target dose of 100 mg/m² and compared to Actellic® 300CS, a WHO/PQ-listed organophosphate IRS insecticide applied at recommended label rate of 1000 mg/m^2^ as a positive control. IRS treatments were randomly allocated to the experimental huts. Two replicate huts were used for each VECTRON™ T500 concrete and mud-walled treatment. Two untreated huts (1 concrete walled and 1 mud walled) were used as negative controls. The walls and ceiling of each experimental hut were sprayed using a Hudson X-pert compression sprayer equipped with an 8002 flat-fan nozzle. To improve spraying accuracy, spray swaths were pre-marked on hut walls and a guidance pole was attached to the end of the spray lance to maintain a fixed distance to the wall.

**Table 1:** Experimental hut treatments.

| **N°** | **Treatment** | **Wall Substrate** | **Number of replicate** | **IRS application dose (mg/m<sup>2</sup>)** |
| --- | --- | --- | --- | --- |
| 1 | Control (sprayed with water) | Concrete | 1 | N/A |
| 2 | Control (sprayed with water) | Mud | 1 | N/A |
| 3 | VECTRON™ T500<br>(Broflanilide WP) | Concrete | 2 | 100 |
| 4 | VECTRON™ T500<br>(Broflanilide WP) | Mud | 2 | 100 |
| 5 | Actellic® 300CS<br>(Pirimiphos-methyl CS) | Concrete | 1 | 1000 |
| 6 | Actellic® 300CS<br>(Pirimiphos- methyl CS) | Mud | 1 | 1000 |

### Assessing the quality of spray applications

Before spraying, 5 filter papers (Whatman No.1) measuring 5cm × 5cm were fixed at 5 positions on the hut walls to be sprayed. After spraying, they were removed, left to dry for 1 hour and then wrapped in aluminium foil and stored at 4° C (+/-2°C) in a refrigerator, after which they were shipped to CEM Analytical Services Ltd (CEMAS), UK for chemical analysis to assess the quality of the spray applications. Spray quality was also assessed by measuring the volume of insecticide solution in the spray tank before and after spraying each hut to determine the amount of insecticide solution applied in each hut and the deviation from the predetermined target.

### Hut trial design

Eight (8) consenting human volunteers were selected and trained to sleep in the huts from dusk to dawn (21:00-6:00) for 6 nights each week for a total of 12 months to attract wild free-flying mosquitoes into the experimental huts. Two backup volunteer sleepers were available to fill in should any sleeper be unavoidably absent. To account for individual attractiveness to mosquitoes, sleepers were rotated daily between the 8 experimental huts using a pre-established Latin Square Design.

Wild vector mosquitoes entering the experimental huts were collected for 6 nights each week for 12 months post-IRS application. On the 7^th^ day of each week, the huts were aired in preparation for the next week. Mosquitoes were collected from the different compartments of the experimental hut (room, veranda trap) using a torch and aspirator and placed in correspondingly labelled plastic cups. They were transferred to the laboratory for morphological identification and scoring of blood-feeding and delayed mortality every 24 hours up to 72 hours post-exposure.

### Outcome measures

The efficacy of each experimental hut treatment was primarily expressed in terms of the following outcome measures: *i)* Mortality which is the proportion of female mosquitoes found dead after 24, 48 and 72 hours, *ii)* Exophily measured as the proportion of mosquitoes caught in the exit traps, *iii)* Blood-feeding rates measured as the proportion of blood-fed mosquitoes.

### Residual activity of insecticide treatments

To assess the residual efficacy of the IRS applications on the treated hut walls, unfed 2–5-day-old susceptible *An. gambiae s*.*s*. (Kisumu) and pyrethroid-resistant *An. gambiae s*.*l*. Covè mosquitoes were exposed in WHO cone bioassays to the treated hut walls 1 week after spraying and subsequently at monthly intervals for up to 12 months and every 2 months thereafter for up to 18 months post spraying. At each time point, a total of 50 mosquitoes were tested per hut in cohorts of 10 mosquitoes per cone on each hut wall/ceiling surface. Mosquitoes were exposed to treated surfaces for 30 min following WHO guidelines [29]. Mortality was recorded every 24 h up to 72 h post-exposure.

### Data analysis

Experimental hut data were double entered into pre-established Excel databases and transferred to Stata 14.1 for analysis. Proportional data (exiting rate, blood-feeding, and mortality) were analysed using mixed effects logistic regression while adjusting for the effects of sleeper attractiveness to mosquitoes and clustering by day. Wild vector mosquito mortality was used to assess the non-inferiority of VECTRON™ T500 to Actellic® 300CS using a non-inferiority margin of 0.7 as defined by the WHO [22]. Cone bioassay mortality was pooled for each treatment and substrate at each time point and compared against an 80% cut-off following WHO guidelines [29].

### Ethical considerations

The study received ethical approval from the Ethics Review Committees of the Ministry of Health in Benin (CNERS No. 39) and the London School of Hygiene and Tropical Medicine (No. 1705). Written informed consent was obtained from all human volunteer sleepers participating in the study. Where necessary, the consent form and information sheet were explained to them in their local language. They were also offered a free course of chemoprophylaxis before they participated in the study. A stand-by nurse was available for the duration of the trial to assess any cases of fever or adverse reactions to test items. Any confirmed cases of malaria were treated free of charge at a local health facility.

### Compliance with OECD Good laboratory practice (GLP) principles

Several activities were implemented through the start-up, execution, and reporting of the study to ensure compliance with the OECD principles of GLP. The study protocol was developed by a properly trained study director and approved by the sponsor before starting the study. Equipment used for the study (precision balances for weighing insecticides, the Hudson sprayer for IRS applications in experimental huts, refrigerators for storage of insecticides and filter papers) were calibrated before use. Both IRS products used in the hut trial were verified to be within their expiry dates and were provided with associated certificates of analysis. In addition, the environmental conditions under which these products were stored was verified daily by use of a calibrated data logger. Mosquitoes used for monthly wall cone bioassays were reared and transported in line with established standard operating procedures that ensured the integrity of the strains tested. All computer systems (data loggers, databases, statistical software) used for data collection, entry, and processing, were validated before use. Records were kept of each procedure performed during the study. The quality assurance team of the CREC/LSHTM Facility performed inspections of the study protocol, critical phases of implementation, data quality and final report to assess compliance to GLP and no non-conformances were detected. The final report, along with all study-related documents, are securely stored in the physical and electronic archive of the Facility for up to 10 years. Study inspections performed in 2021 by the South African National Accreditation System (SANAS), the GLP certification body of the Facility, also detected no non-conformances.

## Results

### Susceptibility of wild vector population to broflanilide and other insecticides

Results from the susceptibility bioassays with wild *An. gambiae s*.*l*. Covè vector and *An. gambiae s*.*s*. Kisumu are presented in Fig. 1. Mortality of wild F1 *An. gambiae s*.*l*. reared from larvae collected from breeding sites in the Covè experimental hut station ranged between 6% and 29% with permethrin, deltamethrin and alphacypermethrin thus confirming the high prevalence of pyrethroid resistance in the Covè vector population. *An. gambiae s*.*l*. Covè also showed resistance to DDT (2% mortality), dieldrin (80% mortality) and suspected resistance to bendiocarb (94% mortality). Full susceptibility was observed to broflanilide (100%) and to organophosphates (≥99% with malathion and pirimiphos-methyl).

**Fig. 1.**
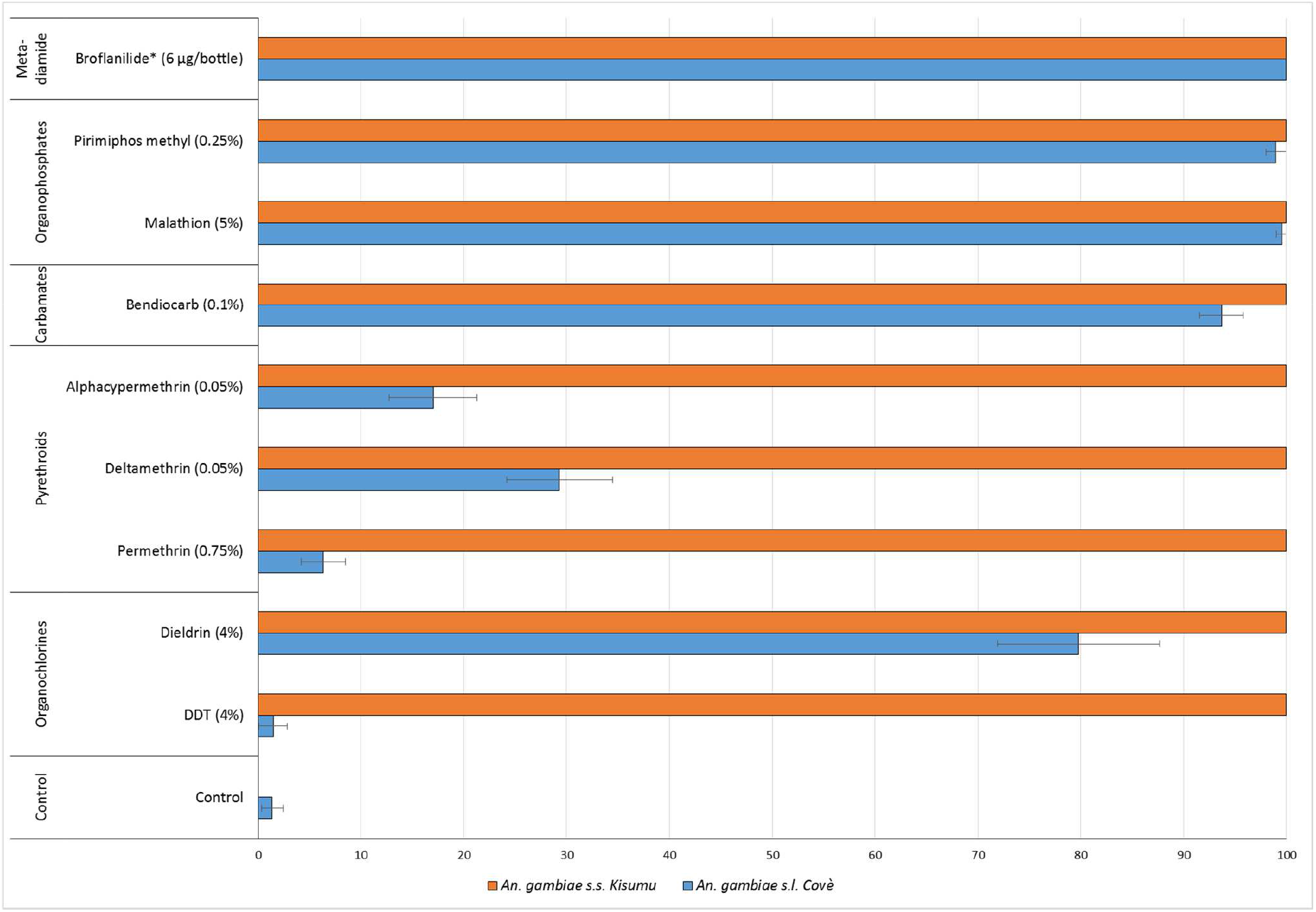
Susceptibility of *An. gambiae s*.*l*. Covè and *An. gambiae s*.*s*. Kisumu to broflanilide and other classes of insecticides used in public health (organochlorine, pyrethroids, carbamates and organophosphates). \**All bioassays were performed in WHO cylinders except for broflanilide, which was tested using the bottle bioassay method*.

Mortality rates with the *An. gambiae* Kisumu strain maintained at the CREC/LSHTM insectary was 100% with all insecticides tested demonstrating full susceptibility of the strain.

### IRS spray quality assessment

A total of 6 experimental huts were sprayed with insecticide while the 2 control huts were sprayed with water only. The percentage deviation from the target insecticide solution volume required for each treatment and the results of the chemical content of filter papers are presented in Fig. 2 and Table 2 below respectively. All spray volumes were within 2% and 18% positive deviation from the target in the treated huts, thus falling within an acceptable 30% deviation (Fig. 2) hence the data showed that the treatments were correctly applied to the experimental huts. This was corroborated by the results of the chemical analysis of filter papers placed on the hut walls before the IRS application. Average AI contents in filter papers showed that VECTRON™ T500 was applied in the range of 51-64 mg AI/m² while Actellic® 300CS was applied in the range of 786-890 mg AI/m². The results from the chemical analysis of filter papers, therefore, showed that all applications were within the WHO-indicated limit of ±50% deviation from the target dose (Table 2) showing that the treatments were correctly applied [31].

**Table 2:** Summary data from chemical analysis of filter papers and assessment of deviation from target dose per treatment. *5 filter papers (Whatman No*.*1) measuring 5cm × 5cm were fixed at 5 positions per huts, sprayed, left to dry and shipped within 2 weeks after IRS application to CEMAS for chemical analysis*.

|  | Control |  | VECTRON™ T500 (100 mg/m²) |  |  |  | Actellic® 300CS (1000 mg/m²) |  |
| --- | --- | --- | --- | --- | --- | --- | --- | --- |
| Wall substrate | Cement | Mud | Cement |  | Mud |  | Cement | Mud |
|  |  |  | Replicate 1 | Replicate 2 | Replicate 1 | Replicate 2 |  |  |
| Target dose (mg/m²) | N/A |  | 100 | 100 | 100 | 100 | 1000 | 1000 |
| Average (mg/m²) | - | - | 61 | 63 | 64 | 51 | 786 | 890 |
| 95% CI | - | - | 44-78 | 51-75 | 40-88 | 33-69 | 534-1038 | 697-1083 |
| Deviation from target dose (%) | N/A | N/A | - 39 | - 37 | - 36 | - 49 | - 21 | - 11 |

**Fig. 2.**
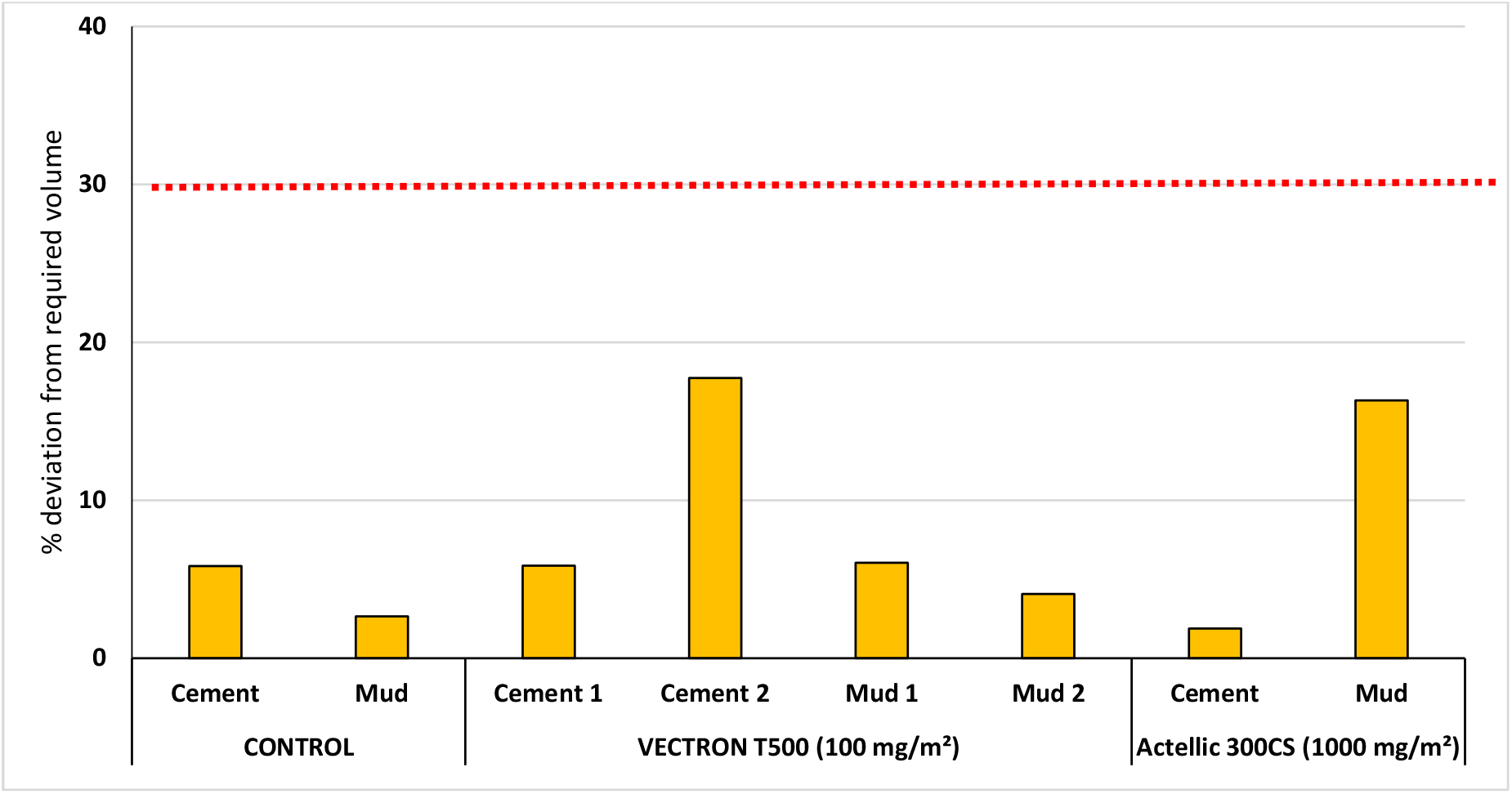
Percentage deviation from the target insecticide volume required for IRS applications in each hut treatment. *The red dotted line indicates the acceptable deviation limit from the target volume (30%)*.

### Mosquito entry and exiting rates of wild pyrethroid-resistant *An. gambiae s*.*l*

The summary of entry and exiting rates of wild, free-flying pyrethroid-resistant *An. gambiae s*.*l*. in each replicate experimental hut is presented in Table 3. Exophily rates for each treatment with pooled data from replicate hut treatments are presented in Fig. 3 below. A total of 23,171 female *An. gambiae s*.*l*. was collected in the experimental over the first 12 months of the trial with the number of mosquitoes caught per hut treatment ranging between 1,696 and 4,627. Both VECTRON™ T500 and Actellic® 300CS showed a deterrent effect relative to untreated huts in both concrete and mud-walled huts though this was higher with VECTRON™ T500 (51%-63%) compared to Actellic® 300CS (23%-45%) (Table 3).

**Table 3:** Entry and exiting rates of wild free-flying pyrethroid-resistant *An. gambiae* s.l. in experimental huts in Covè, Benin.

| Treatment | Control |  | VECTRON™ T500<br>(100 mg/m<sup>2</sup>) |  |  |  | Actellic® 300CS<br>(1000 mg/m<sup>2</sup>) |  |
| --- | --- | --- | --- | --- | --- | --- | --- | --- |
|  |  |  | Cement |  | Mud |  | Cement | Mud |
| Substrate | Cement | Mud | Replicate<br>1 | Replicate<br>2 | Replicate<br>1 | Replicate<br>2 |  |  |
| **Total female collected** | 4 627 | 4 462 | 2 285 | 1 696 | 2 148 | 1 969 | 2 540 | 3 444 |
| **Average per night<sup>α</sup>** | 15 | 14 | 7 | 5 | 7 | 6 | 8 | 11 |
| **% deterrence** | - | - | 51 | 63 | 52 | 56 | 45 | 23 |
| **Total exiting** | 2 000 | 1 817 | 1 657 | 1 399 | 1 561 | 1 469 | 1 938 | 2 517 |
| **% Exophily\*** | 43<sup>a</sup> | 41<sup>b</sup> | 73<sup>c</sup> | 82<sup>d</sup> | 73<sup>c</sup> | 75<sup>c,e</sup> | 76<sup>e</sup> | 73<sup>c</sup> |
| **95% CI** | 41-45 | 38-43 | 70-75 | 80-84 | 70-75 | 72-77 | 74-78 | 71-75 |

<sup>α</sup> The average was obtained by dividing the total female collected by the total number of collection nights (312)
*\*Values in the same row bearing the same letter do not differ significantly at the 5% level according to logistic regression analysis*

**Fig. 3.**
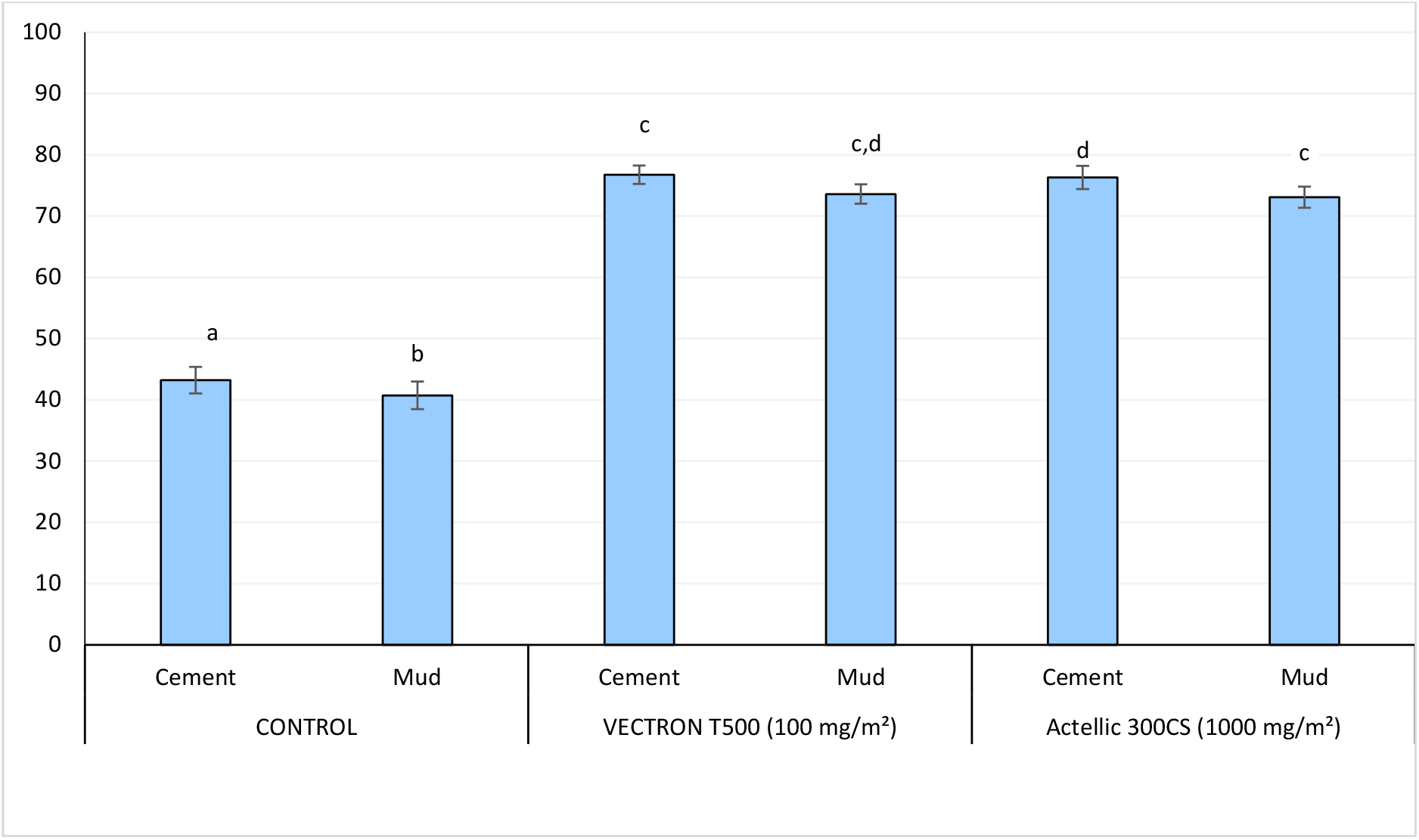
Exiting rates of wild free-flying pyrethroid-resistant *An. gambiae* s.l. in experimental huts in Covè, Southern Benin. *Data for VECTRON*^*TM*^ *T500 cement and mud-walled huts are pooled for both replicates. Bars bearing the same letter label are not significantly different at the 5% level; P>0*.*05, logistic regression analysis. Error bars represent 95% Confidence Intervals*.

Exiting rates with untreated concrete-walled and mud-walled huts (43% and 41% respectively) were significantly lower than what was observed in treated huts (73%-77%, p < 0.001) (Fig. 3). Exophily with VECTRON™ T500 was generally similar to Actellic® 300CS (73%-82% *vs* 73%-76%, p > 0.05) (Table 3).

### Mortality rates of wild free-flying pyrethroid-resistant *An. gambiae s*.*l*. Covè

A summary of overall mortality rates of wild, free-flying, pyrethroid-resistant *An. gambiae s*.*l*. in each replicate experimental hut treatment are presented in Table 4 while mortality results pooled for each treatment are presented in Fig. 4. Over the first 12 months trial, mortality (72h post-exposure) of wild pyrethroid-resistant *An. gambiae s*.*l*. in both control huts was 2% (Table 4 and Fig. 4). VECTRON™ T500 induced 56%-60% overall mortality and this did not differ significantly between both wall substrate-types (56% in cement-walled huts *vs* 60% in mud-walled huts, p = 0.514) while Actellic® 300CS induced 53% mortality in both wall substrate-types huts (Fig. 4).

**Table 4:** Summary overall mortality (24h, 48h, 72h) results of wild, free-flying, pyrethroid-resistant *An. gambiae s.l*. entering experimental huts in Covè, southern Benin for 12 months.

| Treatment |  | Control |  | VECTRON™ T500<br>(100 mg/m²) |  |  |  | Actellic® 300CS<br>(1000 mg/m²) |  |
| --- | --- | --- | --- | --- | --- | --- | --- | --- | --- |
| Substrate |  | Cement | Mud | Cement |  | Mud |  | Cement | Mud |
|  |  |  |  | Replicate<br>1 | Replicate<br>2 | Replicate<br>1 | Replicate<br>2 |  |  |
| Total female collected |  | 4 627 | 4 462 | 2 285 | 1 696 | 2 148 | 1 969 | 2 540 | 3 444 |
| 24h | Total dead | 49 | 37 | 1 021 | 624 | 991 | 864 | 1 296 | 1 772 |
|  | % | 1 | 1 | 45 | 37 | 46 | 44 | 51 | 51 |
|  | 95% CI | 0-4 | 0-4 | 42-48 | 33-41 | 43-49 | 41-47 | 48-54 | 49-54 |
| 48h | Total dead | 60 | 50 | 1 224 | 815 | 1 261 | 1 054 | 1 328 | 1 813 |
|  | % | 1 | 1 | 54 | 48 | 59 | 54 | 52 | 53 |
|  | 95% CI | 0-4 | 0-4 | 51-56 | 45-51 | 56-61 | 51-57 | 50-55 | 50-55 |
| 72h | Total dead | 79 | 72 | 1 330 | 901 | 1 357 | 1 129 | 1 355 | 1 839 |
|  | % | 2<sup>a</sup> | 2<sup>a</sup> | 58<sup>b</sup> | 53<sup>c</sup> | 63<sup>d</sup> | 57<sup>b</sup> | 53<sup>c</sup> | 53<sup>c</sup> |
|  | 95% CI | 0-5 | 0-5 | 56-61 | 50-56 | 61-66 | 54-60 | 51-56 | 51-56 |

*\*Values bearing the same letter superscript along a row are not significantly different at the 5% level (P>0.05)*

**Fig. 4.**
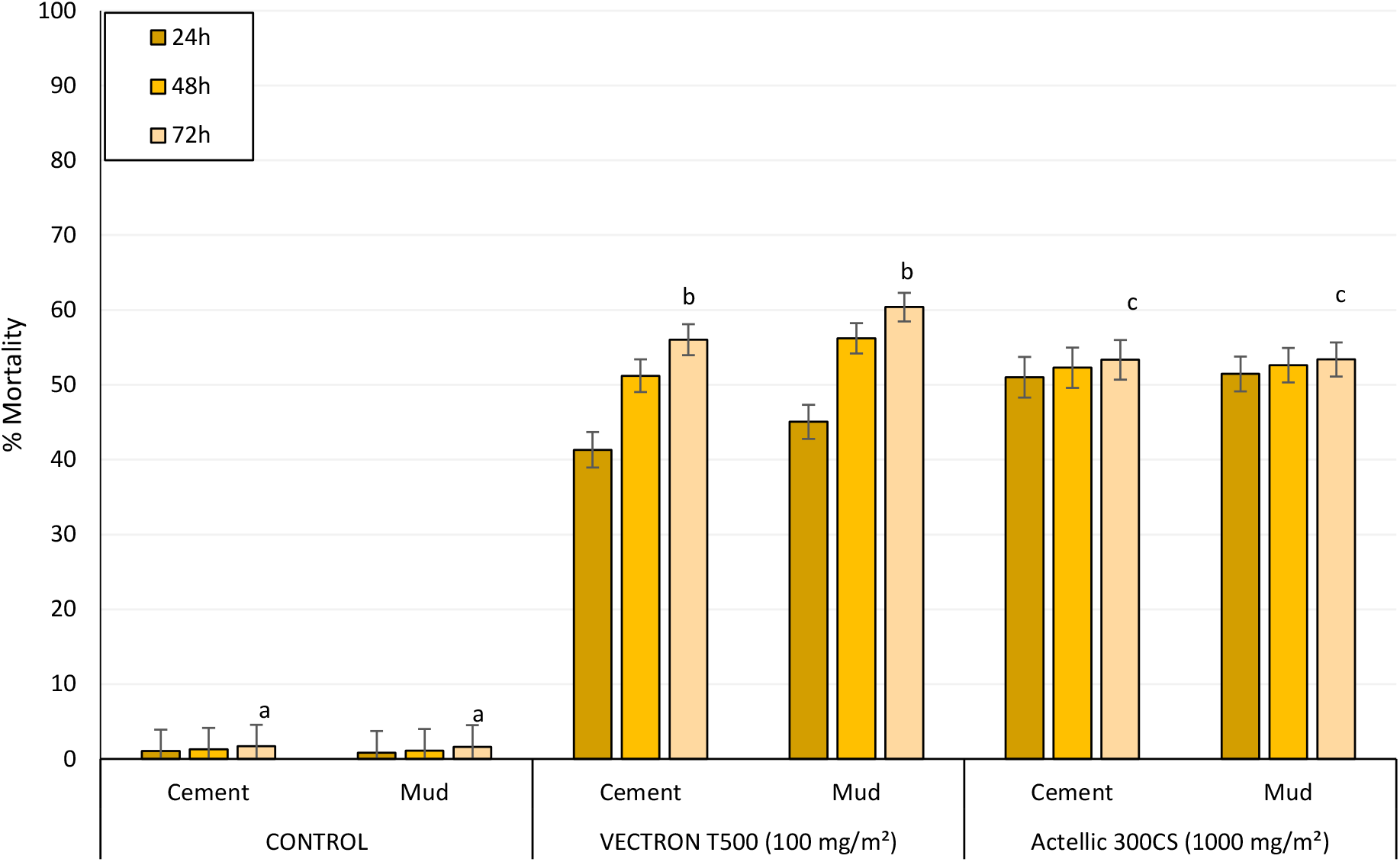
Overall mortality (24h, 48h and 72h) of wild, free-flying, pyrethroid-resistant *An. gambiae s.l*. entering experimental huts in Covè, southern Benin for 12 months. *Overall mortality data for VECTRON*^*TM*^ *T500 cement and mud-walled huts are pooled for both replicates. For 72h mortality, bars bearing the same letter label are not significantly different at the 5% level; P>0*.*05, logistic regression analysis. Error bars represent 95% Confidence Intervals. VECTRON*^*TM*^ *T500 induced a delayed mortality effect*.

VECTRON™ T500 demonstrated a clear delayed expression in mortality irrespective of the type of wall substrate type (Fig. 4). Overall mortality with VECTRON™ T500 increased from 41%-45% at 24h post-exposure to 56%-60% at 72h post-exposure. This delayed mortality effect was not observed with Actellic® 300CS; mortality was 51% at 24h and 53% at 72h post-exposure.

Wild mosquito mortality with VECTRON™ T500 was 62%-73% in the first 3 months and remained >50% for 9 months on both concrete and mud wall substrate meanwhile mortality in huts treated with Actellic® 300CS was 72-95% in the first three months but declined sharply to <40% after 4 months (Fig. 5). Wild vector mosquito mortality in VECTRON™ T500 treated huts was, therefore, moderate but prolonged and steady lasting up to 9 months on both substrate types.

**Fig. 5.**
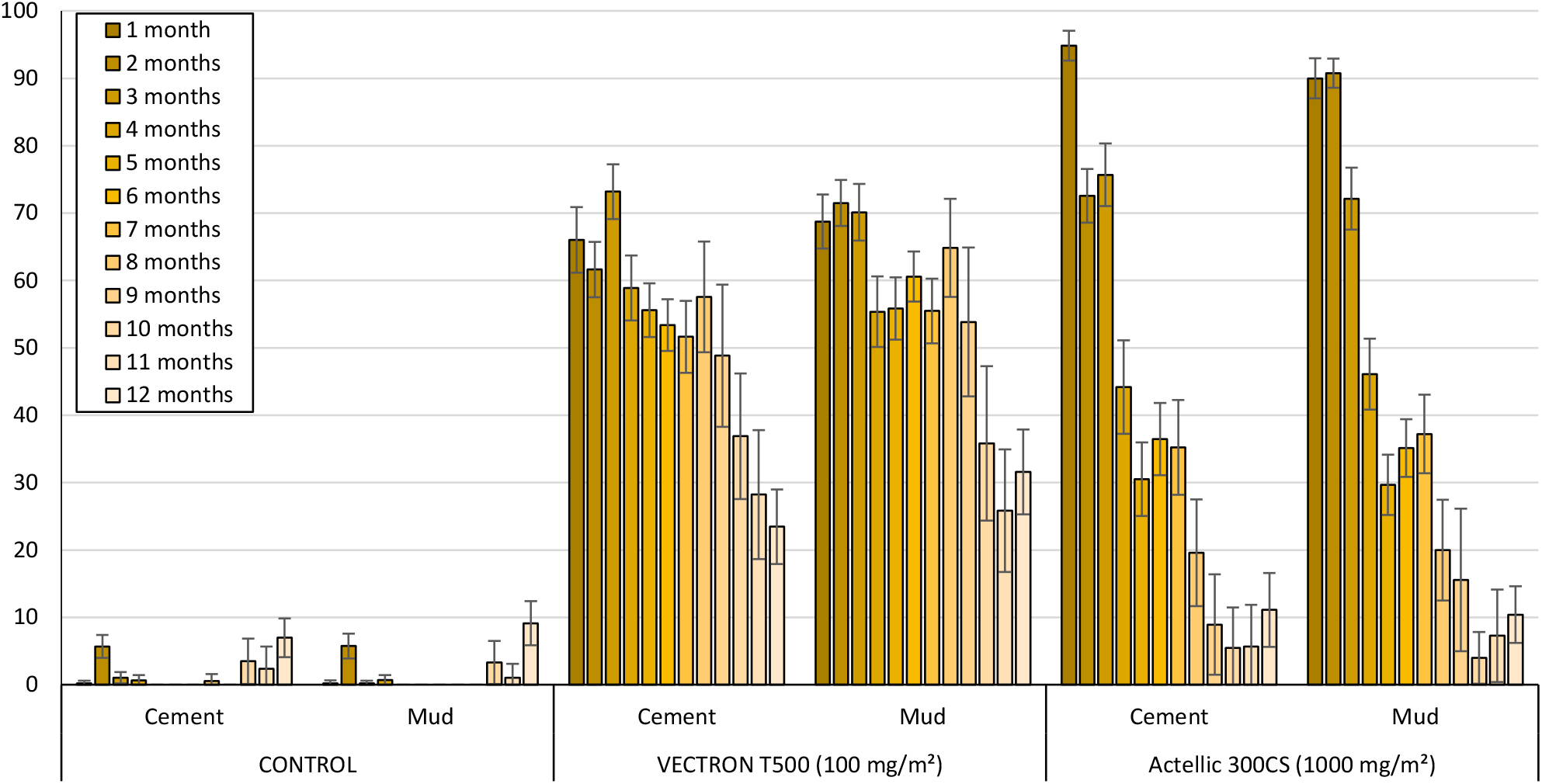
Monthly mortality rates of wild free-flying pyrethroid-resistant *An. gambiae s*.*l*. entering experimental huts in Covè, Southern Benin for 12 months. *Monthly mortality data for VECTRON*^*TM*^ *T500 cement and mud-walled huts are pooled for both replicates*.

### Blood feeding rates of wild pyrethroid-resistant *An. gambiae s*.*l*. Covè

Table 5 shows the results for blood-feeding of wild, free-flying, pyrethroid-resistant *An. gambiae s*.*l*. throughout the trial in each replicate hut while Fig. 6 shows blood-feeding rates pooled per treatment. As expected of IRS treatments, blood-feeding rates were generally very high across all treatments (>96%). Mosquito blood-feeding with the untreated mud and cement huts was 99%. With VECTRON™ T500, the pooled blood-feeding rates were respectively 98% and 97% on mud and cement; and did not differ from what was observed with Actellic® 300CS on either substrate (98%, p > 0.05) (Fig. 5).

**Table 5:**
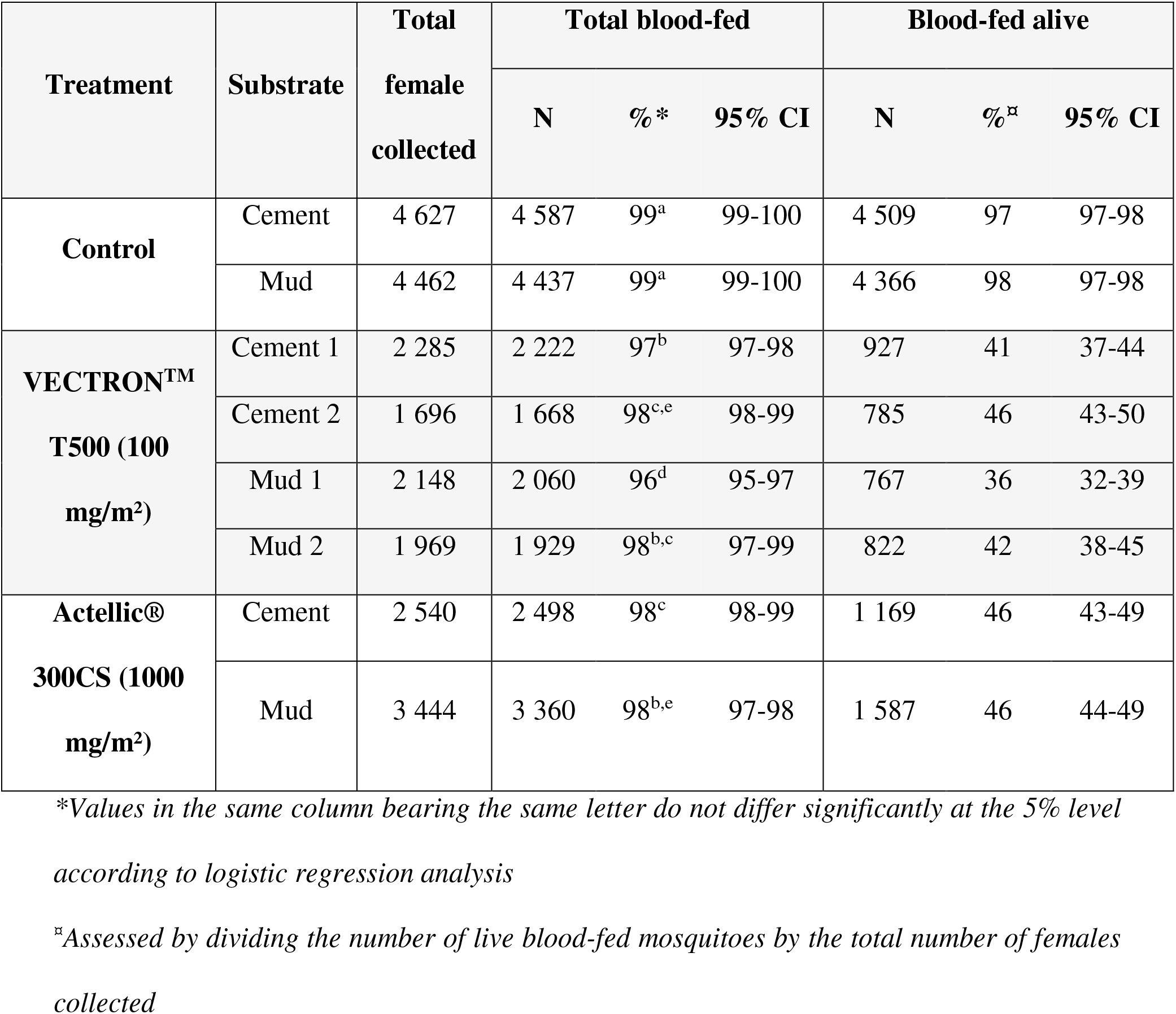
Summary of blood-feeding results of wild, free-flying, pyrethroid-resistant *An. gambiae s*.*l*. entering experimental huts in Covè, southern Benin.

**Fig. 6.**
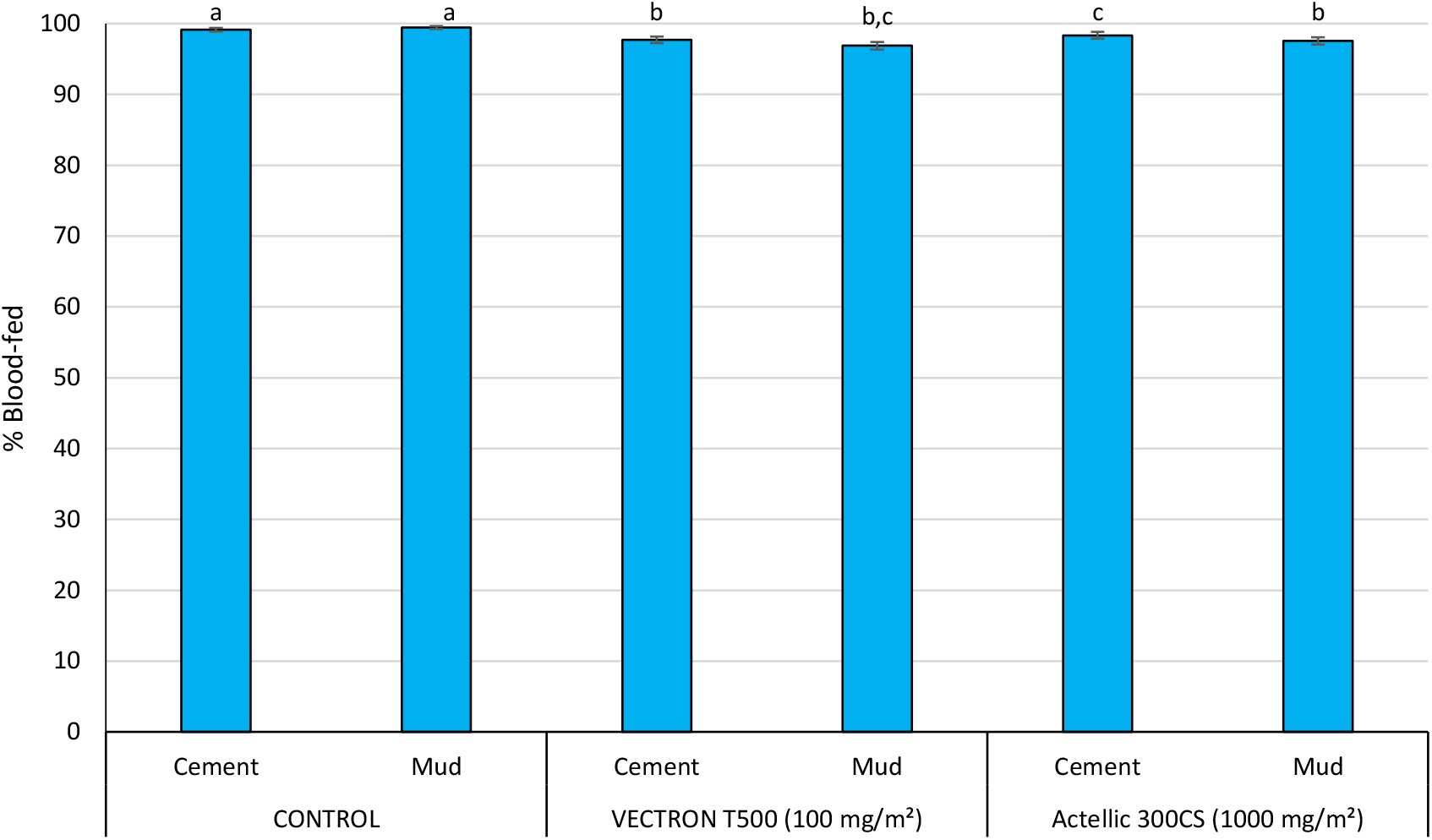
Blood-feeding rates of wild, free-flying, pyrethroid-resistant *An. gambiae s*.*l*. entering experimental huts in Covè, southern Benin. *Blood-feeding data for VECTRON*^*TM*^ *T500 cement and mud-walled huts are pooled for both replicates. Bars bearing the same letter label are not significantly different at the 5% level*.

The proportion of blood-fed mosquitoes that remained alive in the untreated control huts was 97% - 98% of alive *An. gambiae s*.*l*. (Table 5). This proportion reduced to 36%-46% with VECTRON™ T500 and 46% with Actellic® 300CS (46%).

### Residual efficacy of IRS treatments

The results from monthly 30 minutes wall cone bioassays were performed monthly for 18 months after treatment applications using unfed susceptible *An. gambiae s*.*s*. Kisumu and pyrethroid-resistant *An. gambiae s*.*l*. Covè mosquitoes are presented in Figs 7 and 8. With the susceptible Kisumu strain, mortality over 18 months in the untreated huts remained <5%; no correction of mortality data using Abbott’s formula was required. VECTRON™ T500 showed a good residual efficacy on both concrete and mud hut walls as it consistently induced >80% mortality with *An. gambiae* Kisumu for 18 months. With Actellic® 300CS, the mortality of the Kisumu strain remained > 80% for up to 6 and 7 months on concrete and mud wall substrates respectively (Fig. 7).

**Fig. 7:**
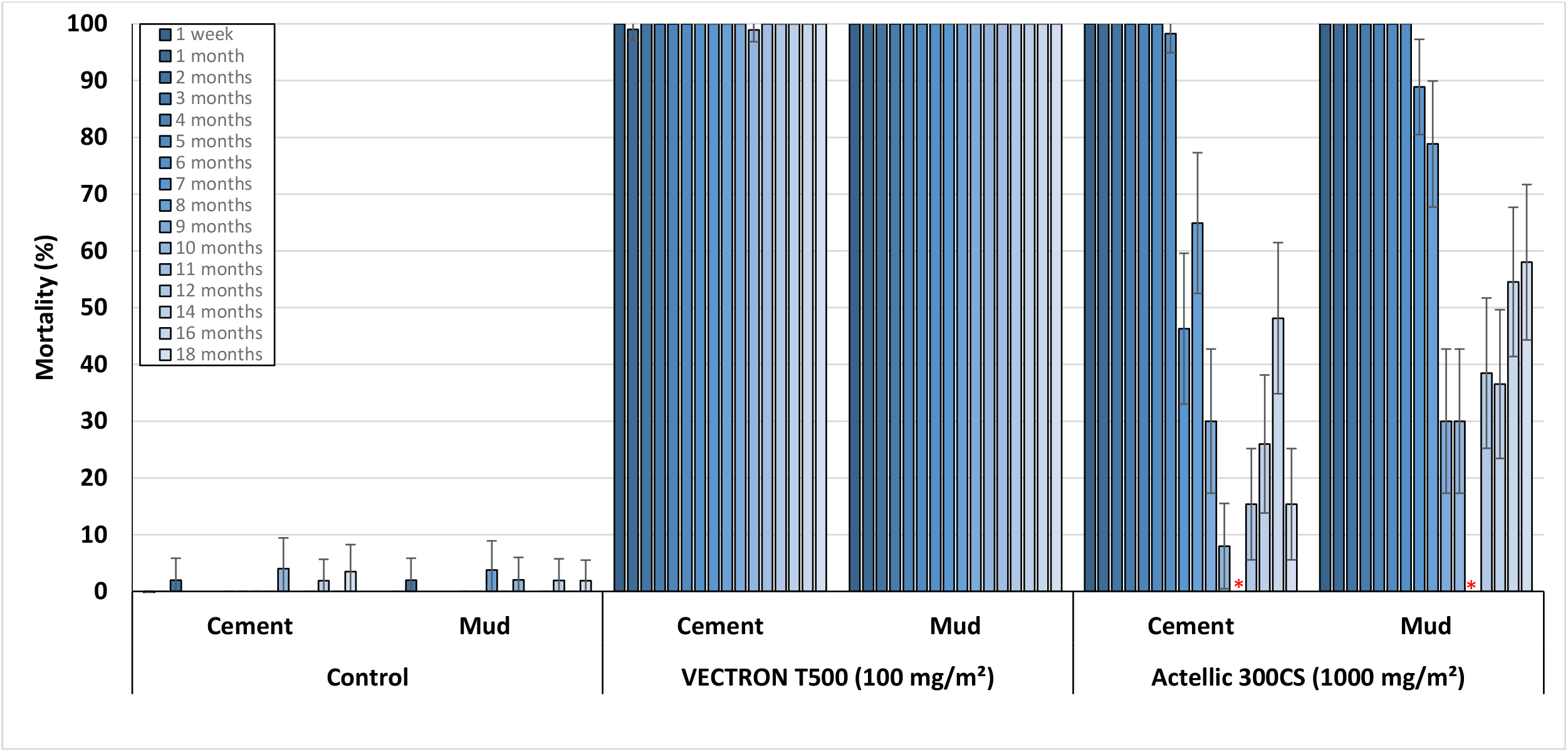
Cone bioassay mortality with susceptible *An. gambiae* s.s. Kisumu on VECTRON™ T500 treated experimental hut walls. *Data for VECTRON*^*TM*^ *T500 treated cement and mud-walled huts are pooled for both replicates*.* Testing not performed at month 11 for Actellic® 300CS due to low mosquito availability

**Fig. 8:**
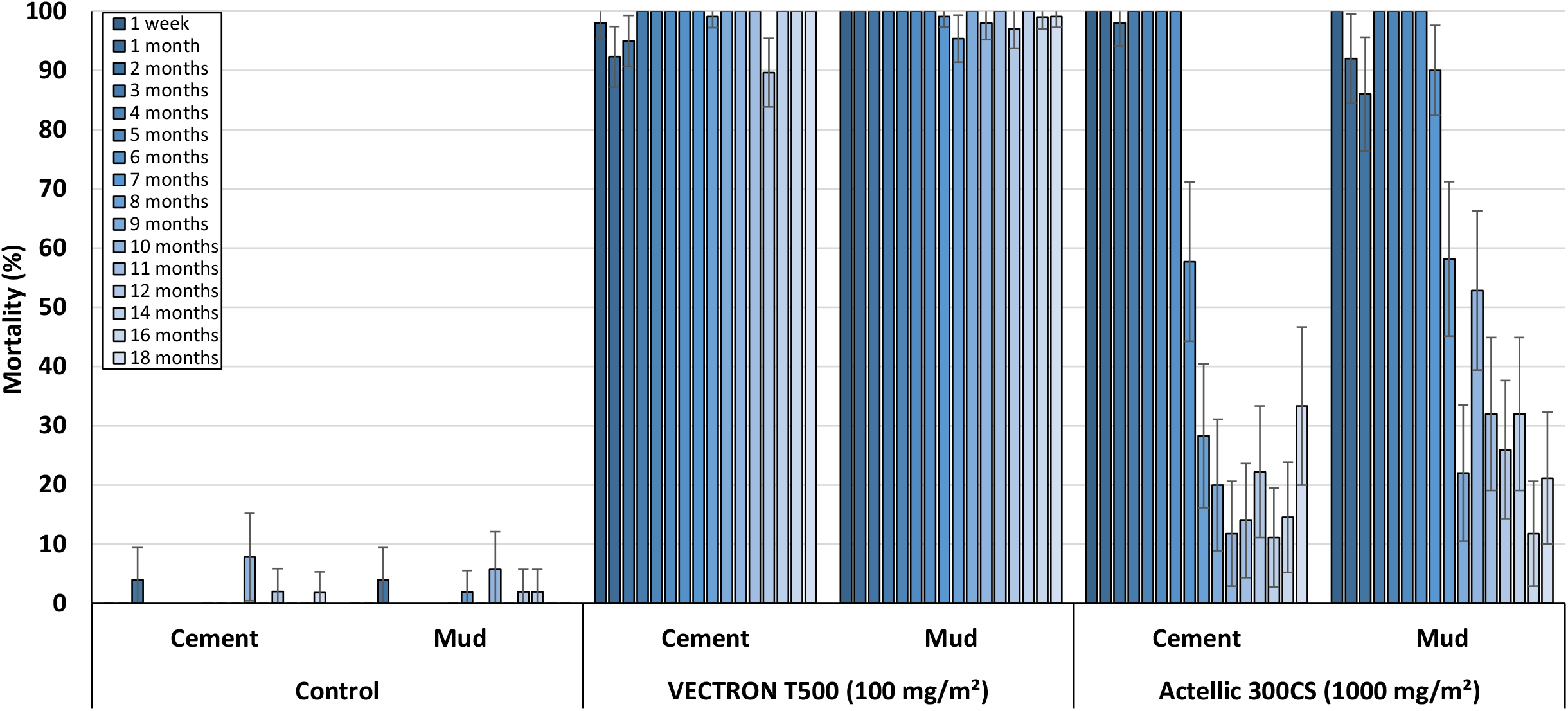
Cone bioassay mortality with pyrethroid-resistant *An. gambiae s*.*l*. Covè on VECTRON™ T500 treated experimental hut walls. *Data for VECTRON*^*TM*^ *T500 treated cement and mud-walled huts is pooled for both replicates*.

The trend was the same in WHO cone bioassays using the pyrethroid-resistant *An. gambiae s*.*l*. from Covè. Mortality in the control huts was <5% while VECTRON™ T500 consistently induced >80% mortality for 18 months on both mud and cement hut walls. By contrast, Actellic® 300CS induced >80% cone bioassay mortality with the pyrethroid-resistant strain of *An. gambiae s*.*l*. from Covè, for 6 months on cement hut walls and 7 months on mud hut walls (Fig. 8).

### WHO non-inferiority assessment

Outcomes of the assessment of non-inferiority between VECTRON™ T500 and Actellic® 300CS are presented in Table 6 below. The odds ratio describing the difference in overall wild mosquito mortality in concrete walled huts between VECTRON™ T500 (56%) and Actellic® 300CS (53%) was 1.240 (95%CI: 1.106-1.391) while the odds ratio describing the difference in wild mosquito mortality in mud-walled huts (60% with VECTRON™ T500 *vs* 53% with Actellic® 300CS) was 1.166 (95%CI: 1.042-1.304). Based on the WHO non-inferiority criteria [31], VECTRON™ T500 was non-inferior to Actellic® 300CS in terms of mortality induced in wild, free-flying pyrethroid-resistant *An. gambiae s*.*l*. entering mud and concrete-walled experimental huts in Covè, Benin.

**Table 6:**
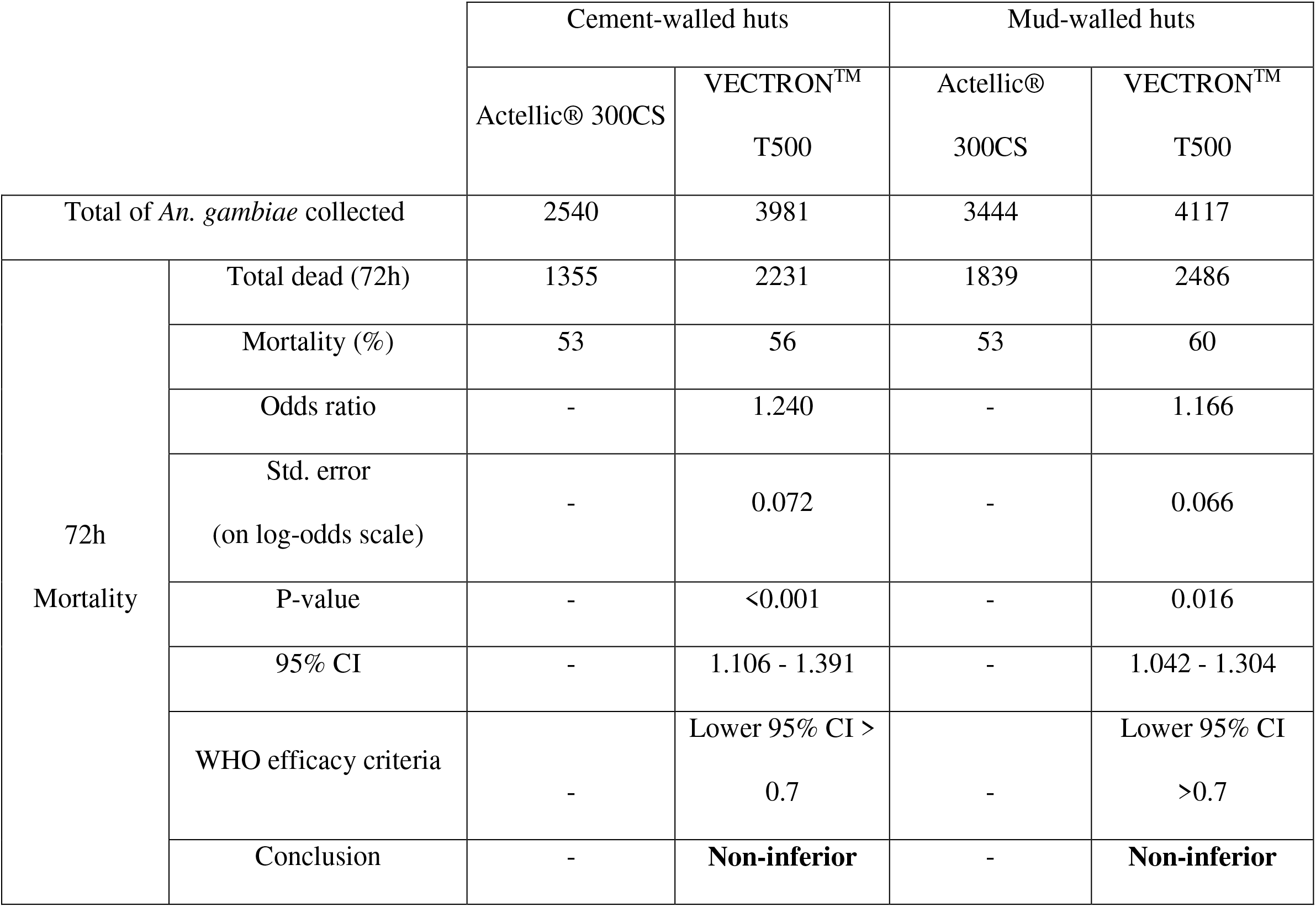
Non-inferiority analysis of mosquito mortality between VECTRON™ T500 and Actellic® 300CS treated cement and mud-walled experimental huts in Cove, Benin.

|  |  | Cement-walled huts |  | Mud-walled huts |  |
| --- | --- | --- | --- | --- | --- |
|  |  | Actellic® 300CS | VECTRON™<br>T500 | Actellic®<br>300CS | VECTRON™<br>T500 |
| Total of *An. gambiae* collected |  | 2540 | 3981 | 3444 | 4117 |
| 72h<br>Mortality | Total dead (72h) | 1355 | 2231 | 1839 | 2486 |
|  | Mortality (%) | 53 | 56 | 53 | 60 |
|  | Odds ratio | - | 1.240 | - | 1.166 |
|  | Std. error<br>(on log-odds scale) | - | 0.072 | - | 0.066 |
|  | P-value | - | <0.001 | - | 0.016 |
|  | 95% CI | - | 1.106 - 1.391 | - | 1.042 - 1.304 |
|  | WHO efficacy criteria | - | Lower 95% CI ><br>0.7 | - | Lower 95% CI<br>>0.7 |
|  | Conclusion | - | **Non-inferior** | - | **Non-inferior** |

## Discussion

Given the current reliance of malaria vector control on insecticide-based strategies, sustained investment is required to develop new active ingredients with novel modes of action [32]. In this study, we investigated the efficacy of VECTRON™ T500, a wettable powder formulation of the newly discovered broflanilide insecticide, for indoor residual spraying. VECTRON™ T500 was evaluated for the first time in human-occupied experimental huts for IRS against a high pyrethroid-resistant malaria vector population in Covè, Southern Benin. At an application rate of 100 mg AI/m^2^ on local wall substrates (cement and mud), VECTRON™ T500 induced prolonged mortality of wild free-flying pyrethroid-resistant malaria vectors entering the experimental huts for 9-10 months. These findings complement previous studies conducted in animal baited experimental huts against wild pyrethroid-resistant *An gambiae* in Covè, southern Benin [23], and pyrethroid-resistant *Anopheles arabiensis* in Moshi in Tanzania [25] that showed the potential of the insecticide to remain efficacious against wild vector mosquitoes for over 6 months. VECTRON™ T500 could, therefore, provide year-round protection to householders in malaria-endemic areas with long-transmission seasons, using a single IRS application round.

To be covered by WHO policy recommendation, a new IRS insecticide must demonstrate non-inferiority to at least one existing IRS product in experimental hut studies [26]. In terms of overall mortality of wild vector mosquitoes, our results demonstrated the non-inferiority of VECTRON™ T500 to Actellic® 300CS, a WHO prequalified organophosphate insecticide formulation that has provided substantial control of mosquito vectors and malaria across distinct ecological zones across Africa [33-37]. VECTRON™ T500, therefore, fulfilled the WHO criteria to be added to the current list of WHO-recommended IRS insecticides in this trial. While it is unclear how residual efficacy in monthly wall cone bioassays relates to the public health impact of an IRS insecticide when applied in communities, the number of months for which an insecticide continues to induce high levels of mosquito mortality (>80%) is used to compare the residual activity of different types of insecticides when applied for IRS [29, 38]. IRS insecticides that show prolonged efficacy in cone bioassays on local wall substrates are in high demand as they can cover the entire malaria transmission seasons without the need for multiple IRS campaigns [38]. Whereas cone bioassay mortality on Actellic® 300CS treated hut walls declined below 80% 6-7 months post-treatment, VECTRON™ T500 remained efficacious in wall cone bioassays for 18 months. This residual efficacy (>18 months) demonstrated by VECTRON™ T500 is longer than that which has been reported in similar studies with other newly developed insecticides for IRS against the same vector population (9-12 months) [39-41]. VECTRON™ T500, therefore, shows potential for use in holo-endemic pyrethroid-resistant areas in Africa usually characterised by prolonged transmission seasons. Its extended residual efficacy may also have positive implications on its cost-effectiveness when applied for IRS. Susceptibility bioassays conducted during the hut trial demonstrated high frequencies of pyrethroid resistance and susceptibility to broflanilide in *An. gambiae s*.*l*. from Covè. These data confirmed the increasing unsuitability of pyrethroids for malaria vector control in Benin [42, 43] and worldwide [8] and demonstrated the suitability of broflanilide for the control of wild insecticide-resistant malaria vector populations. The vector population at Covè was also resistant to dieldrin, an insecticide that also acts on the GABA receptor of mosquitoes [44]. This finding shows the absence of cross-resistance to broflanilide through the mechanism of dieldrin resistance in this vector population confirming previous findings [24]. This could be attributed to the differences in the precise site of action of the two classes of insecticide on the mosquitoes’ GABA receptor as demonstrated in previous studies [19]. As new insecticides are developed for vector control, strategies to preserve the susceptibility of local vectors and extend the useful life of the insecticide need to be considered before large scale deployment [14]. Further studies to develop a validated method for monitoring susceptibility of wild malaria vector populations to broflanilide would be advisable.

According to WHO guidelines, new IRS insecticides that demonstrate efficacy and non-inferiority to existing insecticides in experimental hut trials do not need to undergo epidemiological trials to establish their public health value prior to being recommended for large scale deployment [26, 31]. Nevertheless, studies investigating their impact on malaria transmission indices in large-scale community-randomised controlled trials in endemic communities are useful. Community randomised trials are ongoing in Benin and Tanzania to provide information on the impact of VECTRON™ T500 on entomological indicators of malaria transmission as well as the safety, acceptability to householders and ease of application of the insecticide, when deployed on a large scale.

## Conclusion

VECTRON™ T500 demonstrated prolonged efficacy against wild pyrethroid-resistant mosquito vectors when applied for indoor residual spraying on both cement and mud-walled houses. The insecticide was non-inferior to Actellic® 300CS in terms of its impact on wild vector mosquito mortality and showed extended residual efficacy, lasting over 18 months on both wall substrate types. VECTRON™ T500 shows potential to provide substantial and prolonged control of malaria transmitted by pyrethroid-resistant mosquito vectors and could thus be a crucial addition to the current portfolio of IRS insecticides.

## List of abbreviations

IRS: Indoor Residual Spraying
WHO: World Health Organization
PQ: Prequalification team
GABA: γ -aminobutyric acid
WP: Wettable Powder
CREC: Centre de Recherche Entomologique de Cotonou
LSHTM: London School of Hygiene & Tropical Medicine
Kdr: Knockdown resistance
RDL: Resistant to Dieldrin

## Declarations

### Ethical approval

The trial was approved by the Ethics Review Committees of Ministry of Health Benin (Ethical decision n°26 of 28 June 2019, renewal n°34 of 9 September 2020) and the London School of Hygiene and Tropical Medicine (Ref: 1705). Human volunteer sleepers who slept in the huts to attract mosquitoes gave informed consent before they participated in the study. The study details and consent forms were explained to them in their local language by an interpreter. A stand-by nurse was available to sleepers throughout the study to assess any case of fever and persistent headache. Any sleeper testing positive for malaria was immediately withdrawn from the study and treated effectively in line with local public health practices.

### Consent for publication

Not applicable

### Availability of data and material

The datasets used and/or analysed during the current study are available from the corresponding author on reasonable request.

### Authors’ contributions

RG supervised the hut trial, analysed the data, prepared the graphs and contributed to manuscript preparation. AF and MG performed the hut trial and wall cone bioassays. DT performed the susceptibility bioassays. TS, GS, MR and DN contributed to trial design, supervision, and manuscript review. GP provided administrative and logistics support and contributed to manuscript revision. CN co-designed the study, supervised the project and prepared the final manuscript.

## Acknowledgements

We thank Dr. Kunizo Mori of Mitsui Chemicals Agro, Inc for supplying the insecticide. We also thank the technical staff of CREC/LSHTM (Abibath Odjo, Josias Fagbohoun, Estelle Vigninou, Abel Agbevo etc) for their assistance. Quality assurance of the GLP study was led by Ms. Victoria Ariori. We appreciate Dr. Sarah Rees and Ms. Janneke Snetselaar of IVCC for program support. We are grateful to the rice farmers at Covè for their support in the hut study.

